# Improvements in estimating bioaccumulation metrics in the light of toxicokinetic models and Bayesian inference

**DOI:** 10.1101/2022.01.17.476613

**Authors:** Aude Ratier, Christelle Lopes, Sandrine Charles

## Abstract

The surveillance of chemical substances in the scope of Environmental Risk Assessment (ERA) is classically performed through bio-assays from which data are collected and then analysed and/or modelled. Some analysis are based on the fitting of toxicokinetic (TK) models to assess the bioaccumulation capacity of chemical substances via the estimation of bioaccumulation metrics as required by regulatory documents. Given that bio-assays are particularly expensive and time consuming, it is of crucial importance to deeply benefit from all information contained in the data. By revisiting the calculation of bioaccumulation metrics under a Bayesian framework, this paper suggests changes in the way of characterising the bioaccumulation capacity of chemical substances. For this purpose, a meta-analysis of a data-rich TK database was performed, considering uncertainties around bioaccumulation metrics. Our results were statistically robust enough to suggest an additional criterion to the single median estimate of bioaccumulation metrics to assign a chemical substance to a given bioaccumulation capacity. Our proposal is to use the 75^th^ percentile of the uncertainty interval of the bioaccumulation metrics, which revealed an appropriate complement for the classification of chemical substances (*e.g*., PBT (persistent, bioaccumulative and toxic) and vPvB (very persistent and very bioaccumulative) under the EU chemicals legislation). The 75% quantile proved its efficiency, similarly classifying 90% of the chemical substances as the conventional method.

## Introduction

Chemical substances, present in the environment as a result of human activities, are of extreme concern due to their persistence, their capacity in being accumulated within living organisms and their potential toxicity on the different levels of biological organization all along trophic chains (Popek 2018; Cousins et al. 2019). It is particularly crucial to bring reliable and precise information on the bioaccumulation capacity of the different chemical substances. Indeed, the internalized concentrations by exposed organisms almost exclusively depend on external and internal factors and/or the physico-chemical properties of the substance. This step then conditions the way in which relevant links can be established between the exposure concentrations and the likely damages on life-history traits (Arnot and Gobas 2006; Chojnacka and Mikulewicz 2014; Armitage et al. 2021).

To account for such issues, regulations all around the world established threshold metrics to associate a chemical substance present into an organism with a potential level of bioaccumulation capacity. This allows to identify and manage the risks linked to the chemical substances. In Europe, chemical substances are governed by the REACH regulation, adopted by the European Union to improve the protection of human health and the environment from the risks that can be raised by chemical substances (European Commission 2006). Under REACH regulation, in order to evaluate the bioaccumulation capacity of chemical substances, the calculation of bioaccumulation metrics is required. According to Ratier et al. (2022b), we use the generic expression “bioaccumulation metrics” to denote either bio-concentration factors (BCF) used when exposure is through water, biota-sediment accumulation factors (BSAF) when exposure is through sediment or biomagnification factors (BMF) when exposure is via food. For chemical substances produced or imported between 10 and 100 tonnes per year, bioaccumulation metrics are not mandatory but still required to classify chemical substances as persistent, bioaccumulative and toxic (abbreviated PBT) or as very persistent and very bioaccumulative (abbreviated vPvB). Most of European countries classify chemical substances as bioaccumulative (abbreviated “B”) if the estimated BCF is in [2000; 5000[, or very bioaccumulative (abbreviated “vB”) if it is *>* 5000 (European Commission 2006). Other regulations around the world (Saito et al. 2011; Government Of Canada 1999; United States Environmental Protection Agency 1979) will rather classify as “B” a chemical substance with a bioaccumulation metric ranging in [1000; 5000[. Chemical substances with a bioaccumulation metric in [1000; 2000] are always classified as low bioaccumulative (abbreviated as “𝓁B”), while a bioaccumulation metric *<* 1000 will always correspond to the non bioaccumulative class. These classifications are summarized in Wassenaar et al. (2020) and Hartmann et al. (2014).

Bioaccumulation metrics are calculated from toxicokinetic (TK) parameter estimates by fitting a TK model (usually a one-compartment model) to experimental data collected from bioaccumulation tests (*e*.*g*., OECD (2008, 2012)). Data may be analysed with different tools. Bioaccumulation tests traditionally provide internal concentration measurements from two-phase experiments: a first phase (the “accumulation” phase) during which organisms are exposed via one or several uptake route(s) (water, pore water, sediment and/or food) to a given chemical substance, kept constant over time; a second phase (the “depuration” phase) during which organisms are transferred into a clean medium where elimination processes take place. The TK model is expected to account for all uptake routes and elimination processes (including excretion, dilution by growth and/or metabolization) to properly describe the overall kinetics in terms of internal concentrations. In addition to the experimental procedure of bioaccumulation tests, the OECD guidelines also detail how to obtain bioaccumulation metrics depending on the exposure routes that have been considered within the experiments.

Although several scientific and commercial software environments exist for the calculation of bioaccumulation metrics (*e*.*g*., R, Matlab, Python, Graph-Pad Prism, OpenModel, etc.), few are easy to handle, free and open access with ready-to-use TK modelling tools; namely, with mathematical equations automatically computed from the data entered by the user, and all relevant outcomes provided with a support for their interpretation. To name but a few, there are the Excel macro by Gobas et al. (2020), the “bcmfR” package (OECD 2012) and the MOSAIC_*bioacc*_ web service (http://umr5558-shiny.univ-lyon1.fr/mosaic-bioacc/) that has been recently updated (Ratier et al. 2022b; Charles et al. 2022). Furthermore, if a simple one-compartment TK model (*e*.*g*., *considering only bioaccumulation from one exposure source and excretion as elimination process*) is often sufficient to obtain the bioaccumulation metrics for ERA, the goodness-of-fit can be poor, indicating that such a simple TK model is not fully suitable thus requiring the use of a more complex model. For example, one of the most common complexities to account for is growth of organisms during the bioaccumulation test (OECD 2012). Surprisingly, in such a case, guidelines only recommend to seek advice from bio-statistician and/or pharmaco-kineticist experts. In addition to the fact that there are few user-friendly tools to easily perform TK analyses for non experts, it may be difficult to perform more complex TK modelling (*e*.*g*., accounting for multiple exposure routes, growth dilution and/or biotransformation). To our knowledge, only MOSAIC_*bioacc*_ is today generic enough to analyse any species-compound combination of interest, allowing to account for different exposure routes and several elimination processes simultaneously, automatically adapting the fitted TK model according to the input experimental data. MOSAIC_*bioacc*_ provides all possible bioaccumulation metrics accordingly, namely BCF, BSAF and/or BMF (in kinetic or steady-states versions). Furthermore, benefiting from a Bayesian inference framework, MOSAIC_*bioacc*_ is a convenient tool in support of a facilitated in-depth quantification of the uncertainties delivered as probability distributions. Instead of the classical standard deviation, MOSAIC_*bioacc*_ provides a summary of several useful quantiles. While available on-line since May 2020, the number of users of MOSAIC_*bioacc*_ is continuously growing all over the world, whether they are from academia, regulatory bodies or industry (380 recordings these last 6 months). A publicly available database accompanies the MOSAIC_*bioacc*_ services (http://umr5558-shiny.univ-lyon1.fr/mosaic-bioacc/data/database/index_readme.html), with more than 200 accumulation-depuration data sets collected within published scientific papers (Ratier and Charles 2022). All these data sets were automatically analyzed with MOSAIC_*bioacc*_, full analysis reports being made available via the database directly. Additionally to transparency and reproducibility, all this data can be retrieved for further analyses; this is in line with the FAIR principles, which have become almost mandatory today (Wilkinson et al., 2016).

In ERA, a current major knowledge gap to overcome is the consideration of uncertainty in the modelling approaches used in regulation. Recently, the European Food Safety Authority (EFSA) strongly advocated the need to associate uncertainties with model parameter estimates (EFSA Scientific Committee 2018) in general, emphasizing this need for toxicity indicators in particular (Ockleford et al. 2018). Moreover, recent recommendations have been established to account for uncertainty when using toxicokinetic-toxicodynamic (TKTD) models (Baudrot and Charles 2019). Charles et al. (2021) also highlighted how critical it is to take uncertainty into account when assessing the toxicity of a chemical substance to a range of non-target terrestrial plants by revisiting the species sensitivity distribution (SSD) approach.

Consequently, systematically accounting for the uncertainty around bioaccumulation metrics should be the next step in the improvement of the regulation on classification of chemical substances. Indeed, for the market authorisation dossiers of active chemical substances, the current regulatory guideline asks for single mean or median values as bioaccumulation metrics (European Commission 2013). Despite this approach has its own merits (*e*.*g*., easily deliver a bioaccumulation classification decision value), the question raises what to do if the bioaccumulation metric has a numerical value really close to one of the threshold values from which the classification is decided? Does the precaution principle always apply? A way to overcome this question is to consider uncertainty around the usual point value. However, we identified a lack of efficient and easy tools to handle computer resources, specifically designed to automatically provide uncertainties around any model output, specifically in the field of ecotoxicology. To fill in this gap, as well as to sustain the consideration of uncertainty, it seen urgent to develop ready-to-use tools to provide suitable intervals of possible values in addition to the current practice of bioaccumulation metrics. Subsequent questions then follow about the choice of appropriate summary statistics on the uncertainty the regulation should consider, and how one or the other could change the current classification of chemical substances?

On the basis of the above findings and the identified shortcomings, our paper aims to suggest improvements in the estimation of bioaccumulation metrics. In regards to the quantification of their uncertainty, we worked in the perspective to reinforce the statistical foundations leading to the classification of chemical substances as low-, medium-, very, or non-bioaccumulative. In order to put light for decision makers on a way to handle uncertainty in regulation, we discuss the added value of considering it by exploring different quantiles to classify chemical substances. Serving as a proof-of-concept, our approach could be further extended to include more data sets covering a broader range of species and compound combinations to validate a new suitable decision threshold for ERA.

In the following sections, we first briefly present the last updates recently brought to MOSAIC_*bioacc*_, especially the innovative prediction tool that can be specifically used in designing new experiments in full respect of the 3R principles (Replacement, Reduction and Refinement) ensuring animal welfare and quality of science (Prescott and Lidster 2017; Russell and Burch 1959). Then, the added value of accounting for uncertainty of bioaccumulation metrics is underlined through a meta-analysis of the TK database associated with MOSAIC_*bioacc*_. Finally, we demonstrate how influential may be the consideration of uncertainty when classifying chemical substances according to the current regulatory intervals into which the bioaccumulation metric estimates fall.

### Calculations and predictions of bioaccumulation capacity

This section gives a brief overview of the different features of MOSAIC_*bioacc*_. A focus is first made on the calculation of bioaccumulation metrics. Then the new prediction tool is introduced, to finish this first section with illustrative case studies.

#### Calculations of bioaccumulation metrics

Recent updates have been brought to the MOSAIC_*bioacc*_ web application to increase the speed of calculations and improve its user-friendliness. Above all, a new R-package is today available on the official CRAN web site (https://CRAN.R-project.org/package=rbioacc) that allows to similarly perform all MOSAIC_*bioacc*_ calculations and graphs directly in the R software with ready-to-use dedicated functions (Ratier et al. 2022a). The new version of MOSAIC_*bioacc*_ has entirely been rewritten to reduce the length of the source code and take advantage of this new package. So, MOSAIC_*bioacc*_ is now based on a tabbed presentation that clarifies and facilitates browsing from one step to the next. A special tab gives all bioaccumulation metrics, appropriately calculated according to the input observed data that the user has uploaded. By default, the kinetics BCF, BSAF or BMF values are delivered, displayed via their entire posterior probability distribution then summarized with their median (that is the 50% quantile) and their 95% uncertainty interval (bounded by the 2.5 and 97.5% quantiles). In addition, users can ask for the corresponding steady-state bioaccumulation metrics if they consider it relevant enough according to the duration of the accumulation phase, accompanied with internal concentrations measurements assumed to have reached the expected plateau.

#### Prediction of bioaccumulation metrics to optimize experiments

A new tab has been added to MOSAIC_*bioacc*_, namely a”prediction” tool, that allows interactive simulations of a TK model under a constant or a time-variable exposure profile. The main aim of this prediction tool is to assist experimenters in optimizing the design of new experiments, based on previous TK analyses. Indeed, when studying a new chemical substance and/or a new species, some information need to be known in advance in full respect of scientific ethic in terms of experiment on living organisms and chemical use and recycling. For example, the exposure concentration, the duration of the accumulation and depuration phases as well as the number of time points at which internal concentrations should be primarily measured need to be defined in advance. Then, based on previous TK analyses for given close species-compound combinations, TK parameter estimates can serve to simulate what could be expected when planning additional time points and/or extending the accumulation phase for example. Moreover, benefiting from the Bayesian framework, the uncertainty around parameter estimates can be propagated towards predictions as well as any function of the parameters (Baudrot and Charles 2019).

This is in agreement with the actual policy about reducing animal testing. Indeed, designing new experiments being informed from models in advance may reveal particular helpful, especially when numerous environmentally realistic exposure scenarios have to be investigated on biota. Model predictions and extrapolations to similar chemical or species could replace or at least reduce the animal testing once one experiment is made to estimate parameters under similar conditions. Besides, bioaccumulation metrics can also be predicted in order to characterise potential exposure, without necessarily requiring experiment to obtain them (*e*.*g*., if the substance has similar physico-chemical properties). In the future of ERA, such a framework could potentially also be helpful to investigate mixture effects, that would combine several chemicals with similar physico-chemical properties.

#### Illustrations with case studies

We provide a collection of three case studies as supplementary information (SI, see the .pdf file) to illustrate various situations where the prediction tool can be helpful:

### Case study 1

Plan an experiment for an already studied species exposed to a different but chemically similar compound (*i*.*e*., with close physico-chemical properties), without accounting for the parameter uncertainty;

### Case study 2

Compare several species exposed to a same chemical substance accounting for the uncertainty around parameter estimates coming from a previous TK analysis conducted on a species phylogenetically (or taxonomically) close to the new set of interest. In such a case, the user will need to enter the required input information but also a tabular file with the joint posterior distribution of the parameters, either coming from a previous MOSAIC_*bioacc*_ TK analysis or from its own TK software.

### Case study 3

A prediction for a same species-compound combination but for different exposure scenarios for which the user may have observed data to which simulations can be compared as a validation step of the to be further exploited.

### Matter of uncertainty in estimating bioaccumulation metrics

From accumulation-depuration data, one output of great interest is the bioaccumulation capacity of a chemical substance which is assessed through the appropriate bioaccumulation metrics according to the exposure route(s) (either the BCF, the BSAF and/or the BMF). So, benefiting from the probability distributions of these metrics (coming from the propagation of the parameter uncertainty itself) is crucial to catch their precision. This latter may indeed influence the classification of the substance as bioaccumulative or not, and if bioaccumulative, influence the choice between “𝓁B”, “B” or “vB” categories, leading to a potential bias in the next steps of ERA if misclassification. In order to illustrate this critical issue for ERA, we present below a meta-analysis of the TK database associated to MOSAIC_*bioacc*_ (Ratier and Charles 2022).

#### The accumulation-depuration TK database

The TK database currently contains 211 accumulation-depuration data sets collected from a literature review and corresponding to a total of 56 studies. The 211 data sets encompass 52 genus, 124 chemical substances, three different exposure routes (water being the main one, sediment and food), 34 data sets with also biotransformation data (that is metabolization data). Figure 1 shows several tree maps performed with the treemap R-package (Tennekes 2017) allowing to visualise the proportion of each chemical category (Figure 1-a) and each genus (Figure 1-b) among the 211 data sets. Pesticides and hydrocarbons are the most represented chemical substances within the TK database, what can be explained by the predominance of data sets on freshwater invertebrates and fish. *Gammarus* and *Daphnia* are the most represented genus, probably due to less ethic exigence with them, while fish studies are rarer. *Gammarus* and *Daphnia* are also the genus for which the most biotransformation data are available.

**Fig. 1.**
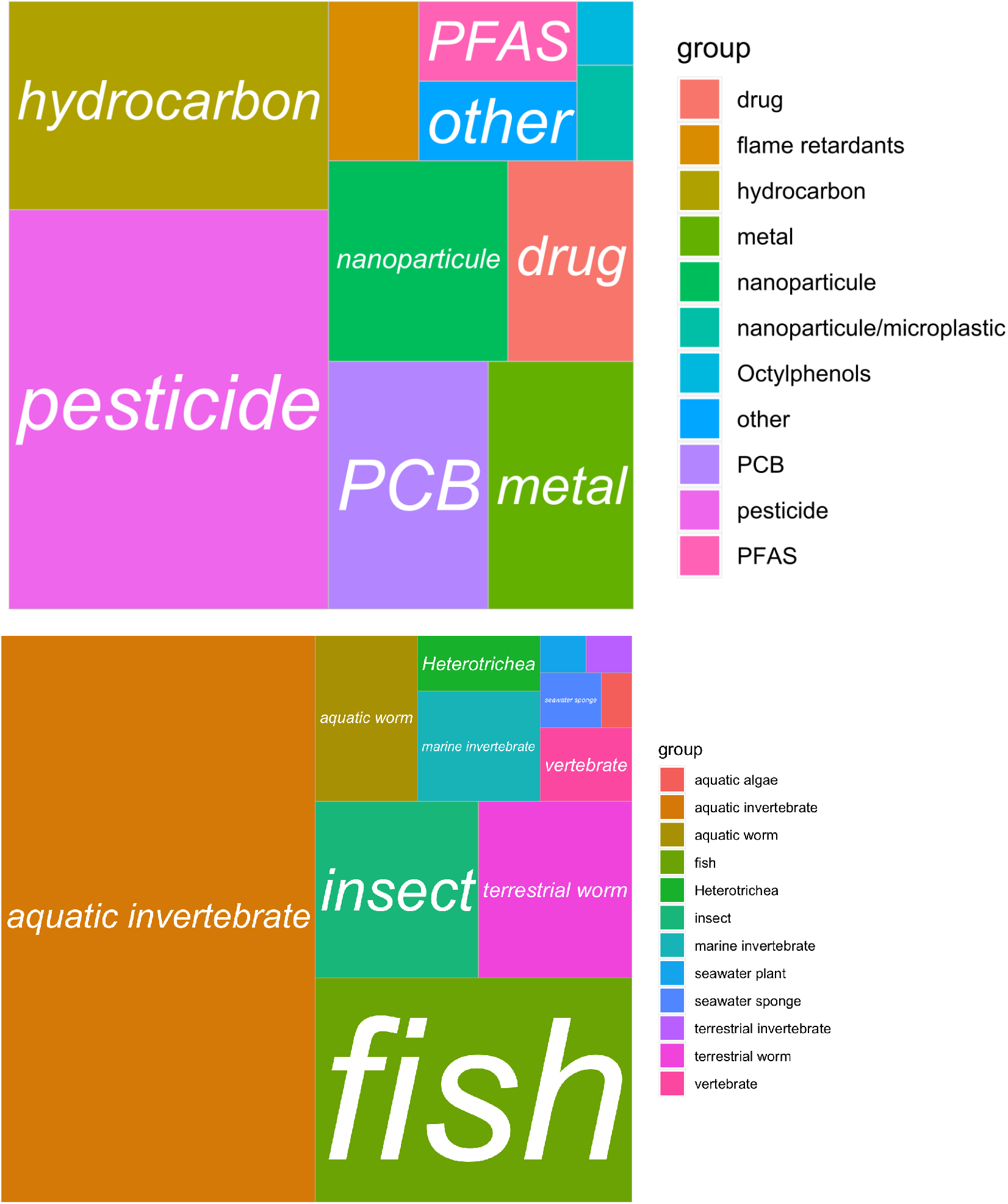
Tree maps of chemical categories (upper panel) and genus categories (lower panel) available in the TK database available at http://lbbe-shiny.univ-lyon1.fr/mosaic-bioacc/data/database/TK_database.html. *Drugs: both human or veterinary pharmaceuticals

The 211 data sets of the TK database were fully analysed with MOSAIC_*bioacc*_. In brief, MOSAIC_*bioacc*_ fits a one-compartment TK model to accumulation-depuration data collected under exposure conditions of interest (Ratier and Charles 2022). MOSAIC_*bioacc*_ automatically builds the appropriate TK model, based on information directly retrieved from the uploaded user data under a predefined format. Accordingly, column headings are assigned to the corresponding model variables, from the subsequent detection of the exposure routes and elimination processes considered in the experimental design. MOSAIC_*bioacc*_ then simultaneously estimates all the model parameters under a Bayesian inference framework. In the end, the posterior probability distributions of bioaccumulation metrics are calculated from the kinetic rate estimates. Based on our experience, for a proper use of MOSAIC_*bioacc*_, we recommend at least six internal concentration measurements during the accumulation phase to ensure the underlying algorithm to converge, whatever the chosen time points, the total duration of the experiment having no matter. Full analysis reports are available from the TK database itself for each data set, as well as references from which data have been extracted. From all these results, we conducted a meta-analysis of the bioaccumulation metrics to specifically illustrate how much accounting for uncertainty matters when characterizing the bioaccumulation capacity of chemical substances.

#### Meta-analysis of the TK database

Even if not required within regulatory documents, associate the uncertainty to a bioaccumulation metric is of crucial importance (Wassenaar et al. 2020). Among the 211 data sets, a total of 137 corresponds to an exposure through water, meaning that MOSAIC_*bioacc*_ analyses 137 BCF probability distributions. Based on the median and the 95% uncertainty interval of these BCF estimates (Figure 2), it appears that aquatic invertebrates have the highest values, when predominantly exposed to pesticides or metals, with *Gammarus* the most represented genus among BCF estimates greater than 5000 classifying the corresponding chemical substances as “vB”. The SI (see the .html file) provides an additional figure with all the 211 estimated bioaccumulation metrics, without distinguishing BCF from BSAF and BMF estimates. Note that from Figure 2, we would have concluded to similar trends as the current practice regarding the bioaccumulation capacity. On the other hand, based on the median of the BCF estimates as required by the regulation, only 22.7% of the BCF values classify chemical substances as “B”, that is with BCF medians higher than 1000. However, this is a fact, such a classification does not rely on the precision of the BCF estimates. Then, we investigated additional decision criteria, namely 75% and 97.5% quantiles, directly extracted from the quantification of the uncertainty around the bioaccumulation metric estimates. We especially evaluated how these new criteria could guide in deciding whether a chemical substance is bioaccumulative or not, and, if yes, with which bioaccumulation capacity.

**Fig. 2.**
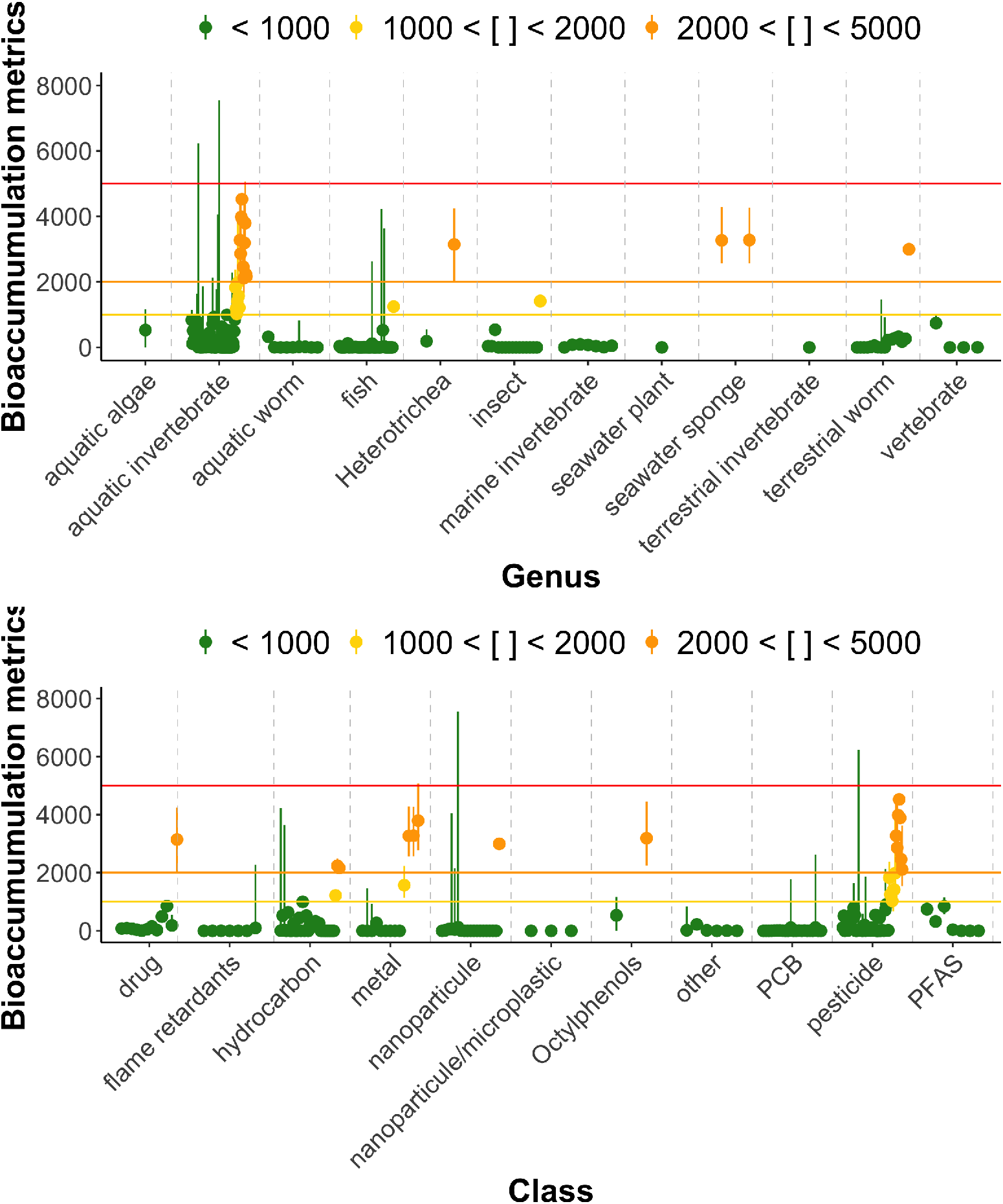
Bioaccumulation metrics according to genus (upper panel) and chemical class (lower panel). Dots represent medians of the bioaccumulation metrics, while vertical segments represent the associated 95% uncertainty intervals. Horizontal lines delineate regulatory threshold values used in the regulation (1000, 2000 and 5000, respectively) to classify chemical substances according to their bioaccumulation capacity. Bioaccumulation metrics are colored accordingly: in green when the metric is *<* 1000; in yellow if the metric is in [1000; 2000[; in orange if the metric is in [2000; 5000[; in red if the metric is *>* 5000.

#### Towards improvements in ERA

ERA can be defined as the evaluation of the incidence and adverse effects likely to occur in the environment given exposure to a chemical substance. ERA is therefore strongly linked to the concept of probability, including both variability and uncertainty of predictions. Accounting for data dispersion allows the quantification of uncertainty, what is today recommended by EFSA (EFSA Scientific Committee (2018)). Although providing central tendency of bioaccumulation metrics remains the core basis in ERA (*i*.*e*., mean or median value), additional information about the uncertainty around this value provides a better picture of the true expectation. Additionally to the biological variability inherent to the data itself, this uncertainty can be due to a lack of data, to an inappropriate or an insufficient experimental design, or to convergence difficulties for example. Among the variety of summary statistics for the uncertainty, standard deviations and minimum/maximum values are probably the most commonly employed, especially when using a frequentist inference process.

Under a Bayesian inference framework, handling uncertainty is very different, the concept of probability being associated to a “degree of belief” rather than to the frequency an even may occur. Starting from a certain confidence in prior knowledge on model parameters, capturing additional knowledge within the data, provides in the end a posterior probability distribution for all parameter estimates. Relevant quantiles extracted from these distributions then allow to summarize the main characteristics of the bioaccumulation metrics. The 50% quantile is the classical median, while the 2.5 and 97.5% quantiles delimit the 95% credible interval, a range that can immediately be interpreted as all possible numbers among which the true bioaccumulation value has a 95% chance of being found.

From here, given that the precision of the bioaccumulation estimates rely on quantiles extracted from their probability distributions, that can be brought into play to investigate new decision criteria for the classification of chemical substances. Instead of the median only, we first considered the upper bound of the uncertainty interval around the bioaccumulation metric estimates, that is the 97.5% quantiles of the posterior probability distribution (referred as Q97.5 in Table 1). This raised the question of an overestimation of the bioaccumulation capacity of the chemical substances. Indeed, the number of BCF values doubled (from 22 to 45) in the “vB” category (Table 1). So, such a criterion would classify 36.5% of the chemical substances as bioaccumulative (*n* = 77), against only 22.7% (*n* = 48) with the usual median (referred as Q50 in Table 1). In terms of margin of safety, this remains within the regulatory context of ERA. However, reasoning on a single value of the bioaccumulation metric (the median or even the mean) may induce a potential misclassification of the substance and, consequently, an underestimation of the potential risk for biota.

**Table 1.** Number of chemical substances in each bioaccumulation capacity class according to the three decision criteria built on the 50%, the 75% or the 97.5% quantiles of the posterior probability distributions of the bioaccumulation metric estimates: Q50, Q75 and Q97.5, respectively. Abbreviation .𝓁B stands for low bioaccumulative, B for bioaccumulative and vB for very bioaccumulative, respectively.

| Criterion | $\ell$B ($BCF \in [1000; 2000[)$ | B ($BCF \in [2000; 5000[)$ | vB ($BCF > 5000$) |
| --- | --- | --- | --- |
| Q50 | 10 | 16 | 22 |
| Q75 | 11 | 20 | 30 |
| Q97.5 | 10 | 22 | 45 |

With the perspective of benefiting from the uncertainty on bioaccumulation metric estimates to suggest a new classification criterion of chemical substances, we compared the use of the 75% quantile (referred as Q75 in Table 1) to both the usual Q50 or the above-mentioned Q97.5, as a possible compromise. As illustrated in Figure 3 (performed with the vioplot R-package by Adler and Kelly 2020, from data on the species *Daphnia magna* exposed to phenanthrene (Wang et al. 2021), the Q50 classifies the chemical substance as “nB” (*BCF <* 1000), while both the Q75 and the Q97.5 criteria classify it as “B” (*BCF >* 1000). This example illustrates that considering the uncertainty would allow to avoid “false negatives”, that is chemical substances classified as non bioaccumulative while they have a 75% chance of being so. In fact, from Figure 3, it can even be stated that this substance has 59.6% to be non bioaccumulative, and correspondingly 40.4% to be, because the quantile associated with the ERA threshold at 1000 is equal to 59.6%. Over the 211 data sets available in the TK database, the Q75 criterion classified 28.9% of the chemical substances as “B” (*n* = 61) against 17.5% for the Q50 criterion (*n* = 37).

**Fig. 3.**
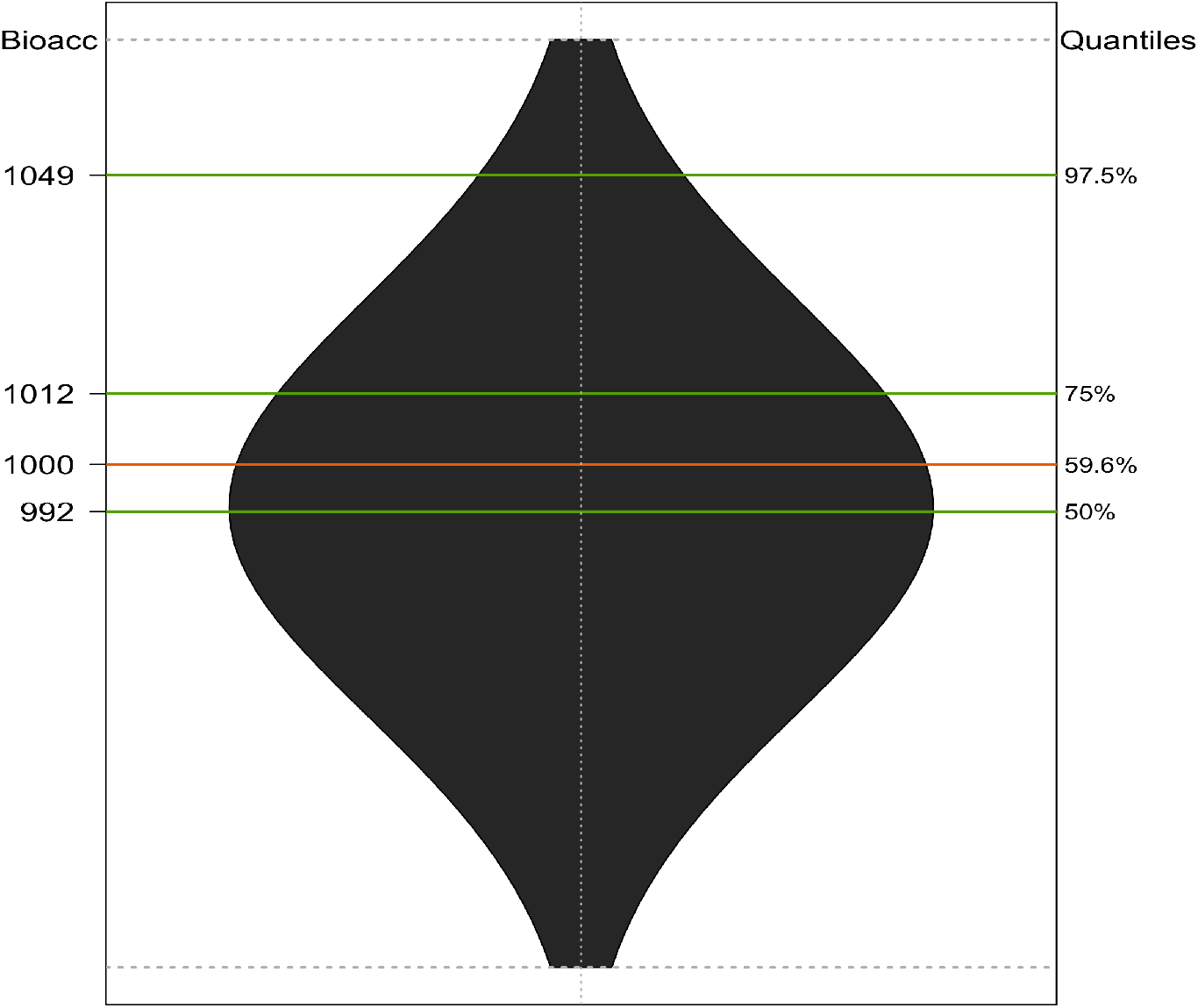
Violin plot of the posterior probability distribution of the bioaccumulation metrics for *Daphnia magna* exposed to phenanthrene (Wang et al. 2021). The left-side scale gives bioaccumulation metric values, while the right-side scale gives the corresponding quantiles of these values. The orange line symbolizes the threshold value at 1000 according to regulatory ERA, while green lines stand for the 50^th^ (Q50 criterion in Table 1), 75^th^ (Q75 criterion) and the 97.5^th^ (Q97.5 criterion) quantiles of the distribution, respectively (Table 1).

Figure 4 shows another example with genus *Enchytraeus* exposed to silver nanoparticles (Topuz and van Gestel 2015), illustrating the possible misclassification of the bioaccumulation capacity of the chemical substance according to the Q97.5 criterion. Indeed, silver nanoparticles are considered as non bioaccumulative (*BCF <* 1000) by both the Q50 (the ERA criterion) and the alternative Q75 criteria, whereas the Q97.5 criterion classified them as “B” (*BCF* ∈ [2000; 5000[). In this case, the difference in the classification comes from a lack of precision of the *BCF* estimate which is associated with a large uncertainty range. This can be assessed with the coefficient of variation (abbreviated as *CV*) which is far over 0.5 in this case. The *CV* is defined in this paper as follows:

**Fig. 4.**
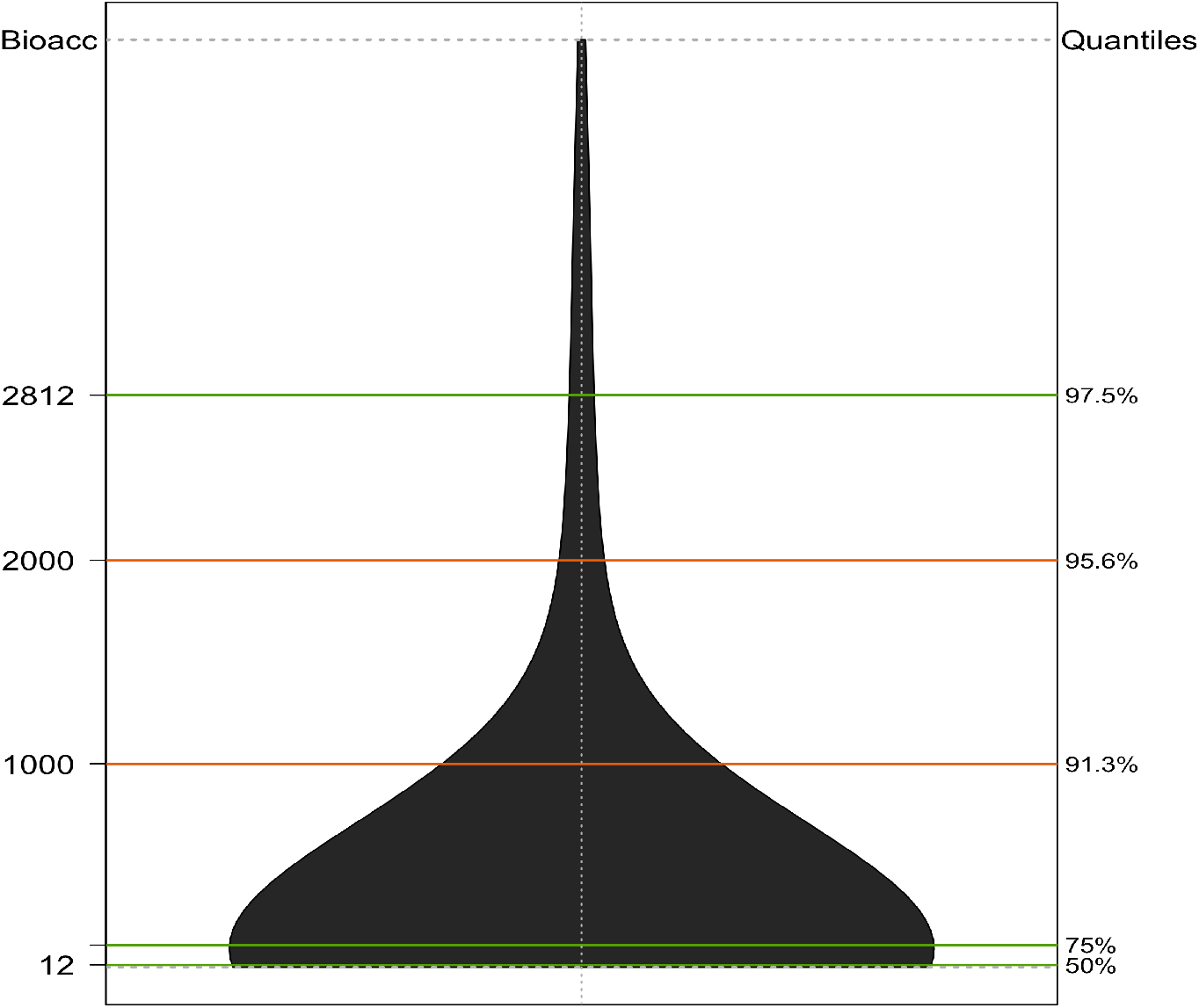
Violin plot of the posterior probability distribution of the bioaccumulation metrics for *Enchytraeus* exposed to silver nanoparticles (Topuz and van Gestel 2015). The left-side scale gives bioaccumulation metric values, while the right-side scale gives the corresponding quantiles of these values. The orange lines symbolize the threshold values at 1000 and 2000 according to regulatory ERA, while the green lines stand for the 50^th^ (Q50 criterion), 75^th^ (Q75 criterion) and the 97.5^th^ (Q97.5 criterion) quantiles of the distribution, respectively (Table 1).

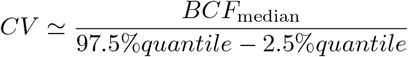

A large uncertainty (*i*.*e*., *CV >* 0.5 of the *BCF* is reflected in Figure 4 by a very dispersed probability distribution towards a highest values (right side of the distribution with a very long tail). On a general point of view, if the estimates of the bioaccumulation metric has a CV *>* 0.5, it will be considered as imprecise. This may be due to a lack of data to first get precise kinetic parameters (*e*.*g*., uptake and elimination rates) or to data that are collected according to an inappropriate experimental design over time. Figure 4 illustrates that imprecise bioaccumulation metrics must be very cautiously interpreted; in this example, the bioaccumulation capacity could have been misinterpreted to decide that the corresponding chemical has a 91.3% chance of being non-bioaccumulative, while the decision threshold is located far in the tail of the distribution, that is within a range of very few probable values.

Considering all the 211 data sets available in the TK database, we performed the classification for all the chemical substances based on each of the three Q50, Q75 and Q97.5 criteria. We then compared the results in order to formulate sufficiently well-founded recommendations and suggest to evolve improve ERA in the future. As illustrated in Figure 5 concerning genus *Anax* exposed to chlorpyriphos (Rubach et al. 2010), the classification was the same whatever the criteria. In total, we assigned more than 90% (*n* = 190) of the chemical substances in to the same class as obtained with the classical Q50 criterion (Table 1). From these results, we finally suggest the use of the 75% quantile of the posterior probability distribution of the bioaccumulation metric estimates as a good compromise to be used complementary to the Q50 and instead of the Q97.5 we also tried. As a consequence, the Q75 could be added to the regulatory ERA requirements to further support decision makers in assessing the bioaccumulation capacity of chemical substances. For this purpose, the MOSAIC_*bioacc*_ web tool could easily be updated with an additional tab automatically delivering both Q50 and Q75 criteria together with the corresponding classification. See SI (.html file) for an example of what this might look like. In the same way, the TK database could be amended to also provide this classification in an all-in-one facility for ERA.

**Fig. 5.**
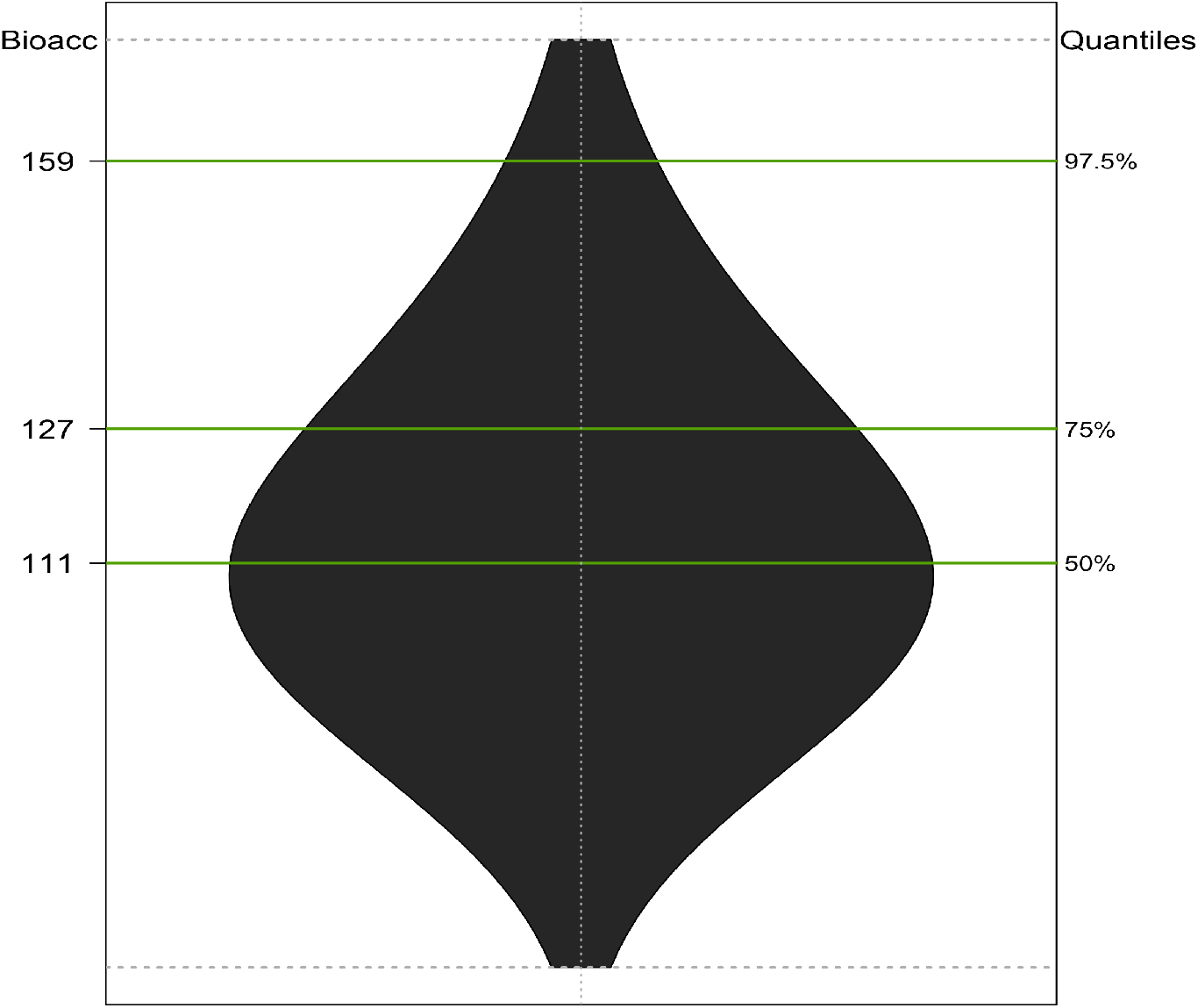
Violin plot of the posterior probability distribution of the bioaccumulation metrics for *Anax* exposed to chlorpyriphos (Rubach et al. 2010). The left-side scale gives bioaccumulation metric values, while the right-side scale gives the corresponding quantiles of these values. The green lines stand for the 50^th^ (Q50 criterion in table 1), 75^th^ (Q75 criterion) and the 97.5^th^ (Q97.5 criterion) quantiles of the distribution, respectively (Table 1).

Over the whole database, using the suggested Q75 criterion, we finally classified 5.2% (*n* = 11), 9.5% (*n* = 20) and 14.2% (*n* = 30) of the chemical substances as “𝓁B”, “B” and “vB”, respectively. This confirms 71% of the chemical substances classified as non bioaccumulative. Focusing on the precision of all bioaccumulation metrics, we got a high variability among species and chemical substances (see SI, .html file). Only considering the Q75 criterion, 37.4% (*n* = 79) of the chemical substances were associated with a *CV <* 0.5, with the high est *CV* among the “nB”-classified chemical substances (38.9%, *n* = 82*/*150). This underlines that for a chemical substances as bioaccumulative, whatever the class, the corresponding bioaccumulation metric was precisely estimated in most of the cases.

Collecting additional TK data sets is the core foundation of our TK database (Ratier and Charles, 2022). Indeed, it has been thought and designed to be dynamically incremented so that researchers are increasingly willing to share their data, either publicly via dedicated repositories, or privately directly to us to be made them available in TK database by referring to the corresponding scientific publication. In turn, they would benefit from automatic reports with MOSAIC_*bioacc*_, for example to revisit previous works or to perform experiments and analyses differently in the future.

## Conclusion

Based on the meta-analysis of the TK database, encompassing today 211 data sets, this paper establishes how crucial it can be to consider the uncertainty in the calculation of bioaccumulation metrics when classifying chemical substances into specific bioaccumulative categories. We fitted TK models under a Bayesian framework, that we specifically chose for its ease of use in delivering of bioaccumulation metrics as posterior probability distributions. This paper thus gathers together statistically-founded results towards the recommendation for a new criterion to sustain this classification. It could indeed be suggested to use the 75% (our preferred indicator) or the 97.5% quantiles of the bioaccumulation metric distributions in order to most appropriately guide the classification of chemical substances into the four regulatory ERA categories: non-, low-, medium-, or verybioaccumulative. The current regulation could accordingly be updated by requesting both the median (50%) and the 75% quantiles to inform decisions on the bioaccumulation capacity. Once MOSAIC_*bioacc*_ improved with the appropriate new features, regulators would be better assisted to classify several chemical substances saving time and gaining confidence in advanced modelling tools.

## Acknowledgments

This work was performed using the computing facilities of the CC LBBE/PRABI.

## Consent for Publication

This manuscript has original research that has not been published previously and is not under consideration for publication elsewhere, in whole or in part.

## Author Contributions

All authors contributed to the investigation of the TK database. Raw data collection and first analyses were performed by Aude Ratier and Sandrine Charles. The first draft of the manuscript was written by Aude Ratier and all authors commented on previous versions of the manuscript. All authors read and approved the final manuscript.

## Supplementary Information and data availability

Supplementary information is available at https://zenodo.org/record/6634584. All data used in this paper is downloadable from the MOSAIC_*bioacc*_ web tool, directly from the associated TK database freely accessible at http://lbbe-shiny.univ-lyon1.fr/mosaic-bioacc/data/database/TK_database.html.

