## Supplementary Information for "Improvements in estimating bioaccumulation metrics in the light of toxicokinetic models and Bayesian inference"

**Archives of Environmental Contamination and Toxicology manuscript No.**
(will be inserted by the editor)

### Supplementary Information: Improvements in estimating bioaccumulation metrics in the light of toxicokinetics models and Bayesian inference

**Aude Ratier**[1,2] · **Christelle Lopes**[1] · **Sandrine Charles**[1,*]

the date of receipt and acceptance should be inserted later

#### Case study 1: Plan an experiment for an already studied species exposed to a different but chemically similar compound

As an example in the case of atrazine, the wish of regulators is to find alternative substances in order to decrease the impact on the environment. From a study on *Daphnia magna* exposed for 48 hours at 100 $\mu g.mL^{-1}$ atrazine spiked water (El-Amrani et al., 2012), regulators are looking to terbuthylazine among other active substances to replace the use of atrazine. To do this, the first step is to collect TK analyses on this similar substance for the same species. Thus, regulators check the corresponding boxes according exposure route (for this example water) and elimination process (here excretion only) in the prediction tool, and fill in the field corresponding to the exposure concentration and the accumulation phase duration. Then the TK parameters values reported in this study are added in the prediction tool (i.e., $k_u = 3.5$ and $k_{ee} = 0.25\ h^{-1}$). Then regulators can choose to test different exposure scenarios, as well as several time for the duration of exposure. From this result, regulators can design experiments according to the output and select appropriate sampling time points for future analyses.

1

Université de Lyon, Université Lyon 1, CNRS UMR5558, Laboratoire de Biométrie et Biologie Evolutive, 69100 Villeurbanne, France.

2

Institut National de l'Environnement Industriel et des Risques (INERIS), Parc ALATA BP2, 60550 Verneuil en Halatte, France.

**Case study 2: Compare several species exposed to a same chemical substance accounted for the uncertainty around parameter estimates coming from a previous TK analysis conducted on a species phylogenetically (or taxonomically) close to the new set of interest.**

For this case study, data already exist for *Chironomus tentans* exposed to benzo-a-pyrene during 3 days by spiked sediment. A user wants to perform an other bioaccumulation test with the same chemical according to the European regulation, but for an other close species (e.g., *Chironomus riparius*). The user requires to choose an exposure concentration in the medium to not exceed the established Environmental Quality Standards (EQS) in the biota, especially focusing on the metabolite. As the distributed parameters from a previous experiment on an other species are available, the user requires to upload them from a previous fit directly performed in $\text{MOSAIC}_{\text{bioacc}}$ or by uploading the corresponding data frame. The results are shown in Figure 1. Benzo-a-pyrene has a threshold value of 0.005 $\mu g.g^{-1}$ in the water European regulation for invertebrates, thus the user can indicates this threshold value in the corresponding field. Besides, time points can be arbitrary plotted according to the values enter by the user depending on the potential experimental design that will be performed. As previously mentioned by (Ratier et al., 2021), with an exposure concentration in the sediment at 0.0136 $\mu g.g^{-1}$, the internal concentration of the metabolite is higher than the threshold value. Thus, the user has to refine the exposure concentration to ensure to be below the threshold, which means to exposed *Chironomus riparius* at 0.011 $\mu g.g^{-1}$.

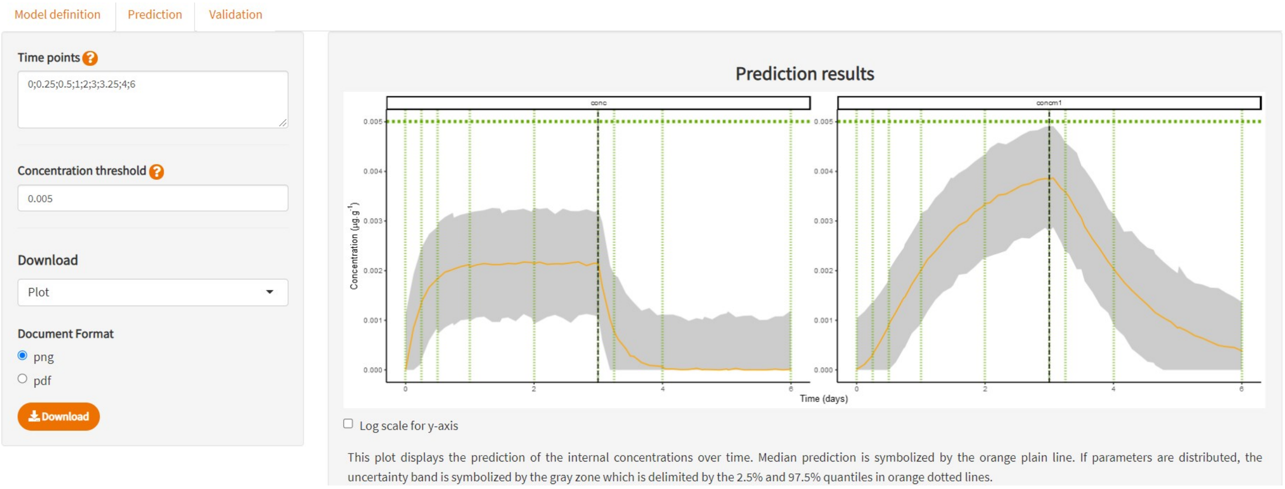


***Fig. 1.*** *Example of predictions from a previous* $\text{MOSAIC}_{\text{bioacc}}$ *analysis data with an other exposure concentration (0.011 instead of 0.0136* $\mu g.g^{-1}$ *initially). The threshold concentration is 0.005* $\mu g.g^{-1}$ *for biota, symbolized by the horizontal dotted green line. Green vertical lines represents the time points for potential samples the user defines by himself.*

**Case study 3: A prediction for a same species/compound combination but different exposure scenarios for which the user may have observed data to which simulations can be compared as a validation step of the exploited TK model**

For a data set with several exposure profiles, the calibration step (i.e., parameters estimation) is performed with $\text{MOSAIC}_{\text{bioacc}}$ for a given concentration. Then the prediction analysis is done for an other exposure concentration for which the user has data. Then for the validation process, the corresponding experimental data for this predicted exposure profile are plotted over predictions. This concept is illustrated with *Spirostomum* exposed to the pharmaceutical product fluoxetine at 0.025 $\mu g.mL^{-1}$ by spiked water for six days for the calibration step (Nalecz-Jawecki et al., 2020). The prediction tool is set up at 0.1 $\mu g.mL^{-1}$ for the exposure phase with identical duration of accumulation and depuration phase. Besides the input parameter are those estimated for the exposure concentration at 0.025 $\mu g.mL^{-1}$. The validation step is performed with measured data from the same study but with an exposure concentration of 0.1 $\mu g.mL^{-1}$ by spiked water and then compared to the predictions previously obtained, as illustrated by Figure 2.

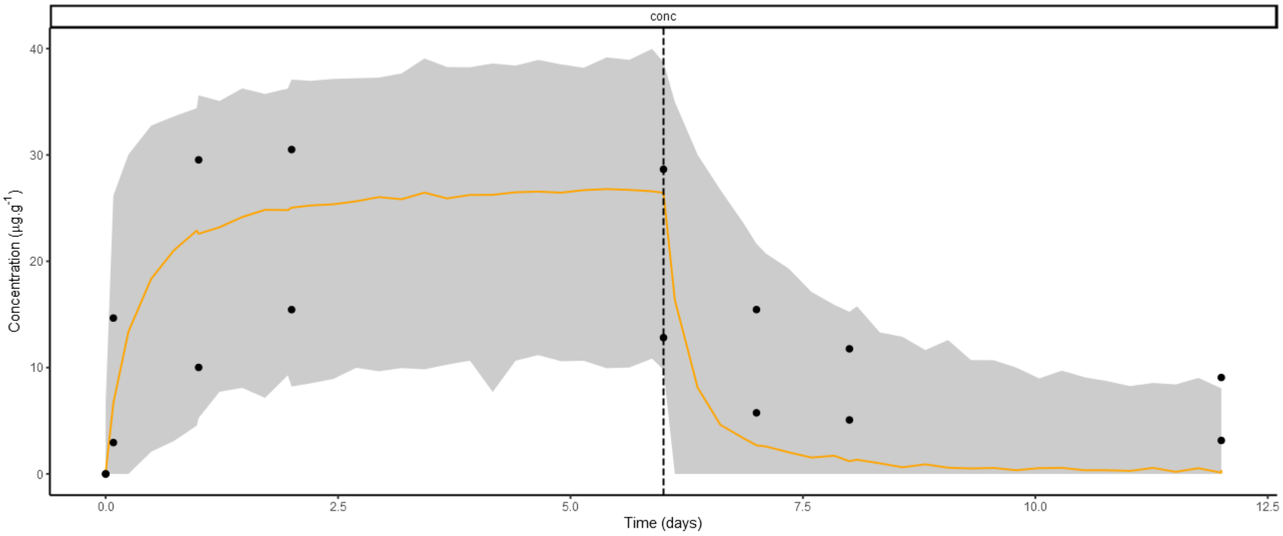


***Fig. 2.*** *Predictions of internal concentrations for Spirostomum exposed to fluoxetine at 0.1 $\mu g.mL^{-1}$ by spiked water for six days. Predictions used distributed parameters from an exposure at 0.0251 $\mu g.mL^{-1}$ citepNalecz-Jawecki2020. Black points represent the validation data (i.e. experimental data for an exposure at 0.1 $\mu g.mL^{-1}$). The orange curve symbolizes median predictions for an exposure at 0.1 $\mu g.mL^{-1}$ bounded by their 95% credible intervals (grey band).*
